# Rapamycin induced autophagy enhances lipid breakdown and ameliorates lipotoxicity in Atlantic salmon cells

**DOI:** 10.1101/2024.12.11.627915

**Authors:** Kanchan Phadwal, Jennifer Haggarty, Dominic Kurian, Judit Aguilar Marti, Ross D. Houston, Mónica B. Betancor, Vicky E. MacRae, Phillip D. Whitfield, Daniel J. Macqueen

## Abstract

Autophagy is a highly conserved cellular recycling process essential for homeostasis in all eukaryotic cells. Lipid accumulation and its regulation by autophagy are key areas of research for understanding metabolic disorders in human and model mammals. However, the role of autophagy in lipid regulation remains poorly characterised in non-model fish species of importance to food production, which could be important for managing health and welfare in aquaculture. Addressing this knowledge gap, we investigate the role of autophagy in lipid regulation using a macrophage-like cell line (SHK-1) from Atlantic salmon (*Salmo salar* L.), the world’s most commercially valuable farmed finfish. Multiple lines of experimental evidence reveal that the autophagic pathway responsible for lipid droplet breakdown is conserved in Atlantic salmon cells. We employed global lipidomics and proteomics analyses on SHK-1 cells subjected to lipid overload, followed by treatment with rapamycin to induce autophagy. This revealed that activating autophagy via rapamycin enhances storage of unsaturated triacylglycerols and suppresses key lipogenic proteins, including fatty acid elongase 6 and acid sphingomyelinase. Moreover, fatty acid elongase 6 was identified as cargo for autophagosomes, suggesting a critical role for autophagy in lipid metabolism in fish. Together, this study establishes a novel model of lipotoxicity and advances understanding of lipid autophagy in fish cells, with significant implications for addressing fish health issues in aquaculture.

## 1. Introduction

Macroautophagy, hereafter referred to as autophagy, is a key cellular recycling mechanism responsible for the degradation of misfolded proteins, damaged organelles, pathogens, and other cellular components in all eukaryotic cells^1^. This process involves the formation of double-membraned autophagosomes, which fuse with lysosomes to form autolysosomes, where the cargo is broken down by lysosomal enzymes and recycled back into the cytoplasm^2^. In addition to the mentioned cargos, lipids stored as lipid droplets (LDs) are degraded through a selective form of autophagy called lipophagy^3^. Excess lipids, if not degraded or stored in LDs, can block autophagic flux, leading to further lipid accumulation and lipotoxicity^4–6^. Lipotoxicity results from the build-up of diacylglycerols and ceramides (CERs) within LDs and other non-adipose tissue^7^. CERs can be further metabolized into complex sphingolipids such as sphingomyelins (SMs), or vice versa^8^. Lipid-induced lipotoxicity is a key contributor to metabolic disorders in mammals, including insulin resistance, myosteatosis, cardiac pathologies, and non-alcoholic fatty liver disease (NAFLD) ^9–11^.

Atlantic salmon (*Salmo salar* L.) is the most commercially valuable farmed fish globally and an important source of high-quality protein and omega-3 fatty acids, supporting food production and representing the most advanced aquaculture sector^12^. Sustainable salmon aquaculture is challenged by a range of issues that are negatively impacting fish health and welfare, including infectious diseases caused by diverse pathogens and parasites^13^. Atlantic salmon health is also thought to be impacted by recent changes in farmed fish diets that are more environmentally sustainable and cost-effective, increasingly replacing protein and oils traditionally derived from marine sources, with those from plants^14^. Vegetable oils have more short chain (< C20) n-6 polyunsaturated fatty acids than fish oil, and reduced n-3 long-chain chain polyunsaturated fatty acids, which changes the n-3/n-6 ratio and promotes adipogenesis by lipid deposition in liver, visceral adipocytes and muscle, while causing lipotoxicity and inflammation^15,16^. In Atlantic salmon, as observed in mice^17,18^, diets enriched in vegetable oils have been shown to increase adiposity^15^, which may lead to lipotoxicity. In addition to having negative impacts on fish health, this is likely to negatively impact human consumers of salmon. In this respect, mice showed an increase in lipotoxic hepatic oxylipins and CERs, as well as increased insulin resistance, after consuming farmed Atlantic salmon fed a vegetable oil-enriched diet^19,20^. Oxylipins are oxidized derivatives of polyunsaturated fatty acids, while CERs are bioactive lipids involved in cell signalling, and both are associated with metabolic disorders, liver disease like NAFLD, and inflammation when present in excessive amounts^21,22^

The role of autophagy in regulating lipid accumulation and breakdown remains unexplored in Atlantic salmon, but represents an important target to better understand and improve the health and welfare of farmed fish. In this study, we used a macrophage-like cell line derived from the head kidney (SHK-1)^23^ as an established *in vitro* model to test the hypothesis that autophagy regulates lipid accumulation and breakdown in Atlantic salmon. We demonstrate that LDs colocalize with key lipophagy markers in Atlantic salmon cells, Moreover, omics provides evidence of lipotoxicity that strongly mirrors human NAFLD in lipid-treated SHK-1 cells; this was ameliorated by autophagy induction, which enhanced unsaturated triacylglycerol (TAG) storage and suppressed lipogenic protein expression. We also show that key lipogenic proteins are cargo for autophagosomes, suggesting that autophagy plays a pivotal role in regulating enzymes involved in altering fatty acid composition and lipid metabolism. The suppression of acid sphingomyelinase activity following autophagy induction further highlights the potential role of autophagy in modulating SM metabolism. Finally, we discuss the implications of our findings for understanding mechanisms of lipid metabolism in farmed fish and the potential to promote health in salmon aquaculture.

## 2. Methods

### Cell Culture and Experimental setup

The SHK-1 cell line (ATCC 97111106)^23^ was cultured in complete L15 media (10% FCS, 5% Penicillin/ Streptomycin and 400μL beta Mercaptoethanol). Cells were incubated in the dark, in a 19°C incubator with CO2. Lipid overload was achieved by treating the cells with 2.5% lipid mixture^24^ (Sigma, L0288), which contains non-animal derived fatty acids (2 μg/mL arachidonic and 10 μg/mL each linoleic, linolenic, myristic, oleic, palmitic and stearic acid), 0.22 mg/mL cholesterol from New Zealand sheep′s wool, 2.2 mg/mL Tween-80, 70 μg/mL tocopherol acetate and 100 mg/mL Pluronic F-68 solubilized in cell culture water. Autophagy was induced by treating SHK-1 cells with 1μM rapamycin (RAPA) (APExBIO, A8167). Autophagic flux was blocked using 10nM Bafilomycin-A (Baf-A) (MERK, B1793), while autophagy was blocked using 500nM MRT68921 (MRT) (APExBIO, B6174). LD staining with (boron-dipyrromethene) BODIPY, LC3 and Perilipin 3 (Plin3) done with cells treated for a period between 24-72 hr, with remaining experiments performed using cells treated for 72 hr. DMSO concentration was kept at 0.01% for all experiments. A schematic diagram summarising the experiments is shown in **Figure 1A**.

**Figure 1.**
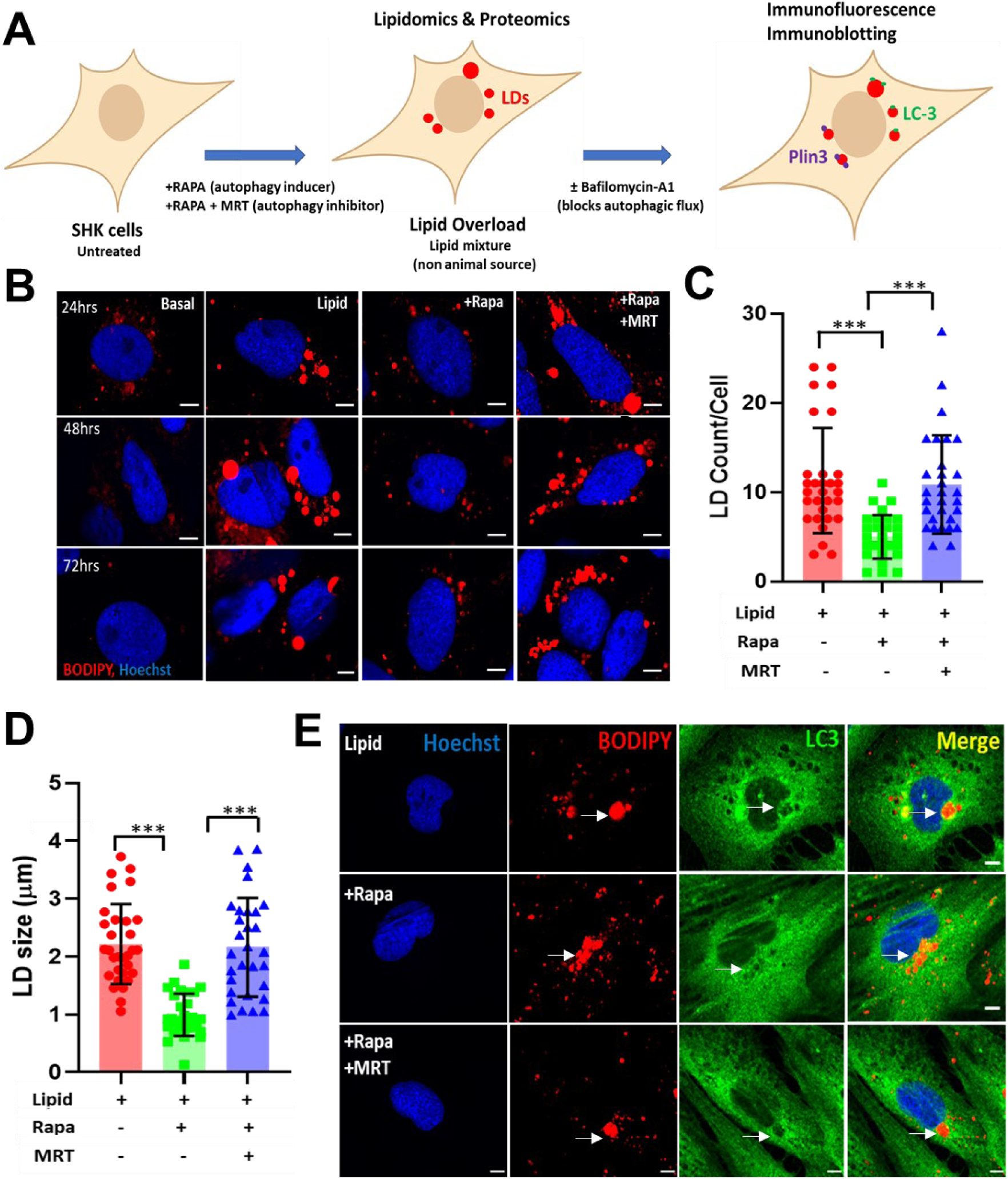
Lipid overload inhibits LD breakdown by blocking autophagic flux in SHK-1 cells. **A.** Schematic overview of experiments performed in this study. **B.** LD count and size measurements in SHK-1 cells following 24, 48 and 72 hr treatments under basal, lipid overload, lipid overload with RAPA, and lipid overload with RAPA+MRT. **C.** Quantification of LD numbers at 48 hr (30 cells counted from 3 independent experiments). **D.** Quantification of LD size at 48 hr (30 cells counted from 3 independent experiments). **E.** Representative confocal images showing the colocalisation of LC3 (green puncta) with LDs stained with BOPIPY (red). Baf-A was added to all treatments to block the autophagic. Scale bar = 10μm, n=3. Data shown as mean +/− SD, *\*\*\*p <* 0.001.

### Lipid droplet and immunofluorescence staining

LDs were stained with 1μM BODIPY 493/503 (D3922). BODIPY 493/503 staining was combined with rabbit polyclonal LC3 (1:300, PM036) and Plin3 (1:300, Abcam, ab47638) for immunofluorescence (IF). Prior to IF, SHK-1 cells were fixed in 4% paraformaldehyde for 20 mins at room temperature and then blocked and permeabilised in blocking buffer (0.2M Glycine, 0.1 mg/mL saponin and 30 mg/mL BSA in PBS) for 1 hr at room temperature. The cells were then incubated with primary antibodies overnight. The cells were then washed three times with PBS and incubated with 1:500 secondary antibody, donkey anti Rabbit 647 (Bio Legend, 406414) for 1 hr at room temperature. After 3 washes, cells were stained with 1:10,000 Hoechst for 1-3 mins and mounted on glass slides with ProLong glass antifade mountant (Invitrogen, P36982).

Using the same methodology, but with an additional blocking step (using 5% goat serum for 45 mins) SHK-1 cells were stained with a combination of 1:250, rat monoclonal LAMP1-546 (SC-19992) and rabbit polyclonal ASM (DF-13384) or rabbit polyclonal ELOVL-6 (DF-4039) on its own. All the experiments were visualised on a LSM880 laser scanning microscope produced (Zeiss) with image analysis performed using Fiji^25^.

### Sphingomyelinase assay

Acid Sphingomyelinase activity was assessed using an Amplite® Fluorimetric Acid Sphingomyelinase Assay *Red Fluorescence* Kit (AAT Bioquest), following the manufacturer’s instructions. SHK-1 cells were cultured in 6 well tissue culture plates and treated for 72 hr (**Figure 1A**). Cells were then washed with cold 2X PBS and scraped into 200μL of RIPA Lysis buffer (Thermo Scientific, 89901). The lysed cell extract was centrifuged at 12,000 RPM for 20 mins at 4°C and the supernatant collected. 50μL each of SM standard, test solution (i.e., cell extract) and a blank solution were added in duplicates to black bottomed 96 well plates. To this, 50μL of SM working solution was added, and plates were incubated for 1 hr at 37°C. Next 50μL of SM working solution was added to each well and further incubated for 1 hr at room temperature. The plates were wrapped in aluminium foil to protect from light. Fluorescence was read at 540/590nm using a BioTek Cytation 3 Cell Imaging MultiMode Microplate Reader.

### Immunoblotting assays

A standard immunoblotting protocol was used for rabbit polyclonal antibodies for LC3 (1:3000, PM036), SQSTM1/p62 (1:500, Ab 264313), ATGL (1:500, Proteintech, 55190-1-AP), and MGL (1:500, Proteintech, 20494-1-AP). Gel quantification was performed using Fiji software. All the blots were normalised against estimated total protein concentration on a per sample basis.

### Lipidomics

Lipids were extracted from SHK-1 cells according to the method described by Folch *et al*^26^. Samples were extracted in a 2/1 (v/v) chloroform/methanol solution. The samples were left at 4°C for 1h then partitioned by addition of 0.1 M KCl. The mixture was centrifuged to facilitate phase separation. The lower chloroform layer was evaporated to dryness under a steady flow of nitrogen gas and reconstituted in methanol containing 5 mM ammonium formate.

Following extraction, lipids were analysed by liquid chromatography-mass spectrometry (LC-MS) using a Thermo Exactive Orbitrap mass spectrometer equipped with a heated electrospray ionization (HESI) probe and interfaced with a Dionex UltiMate 3000 RSLC system (Thermo Fisher Scientific, Hemel Hempstead, UK). Samples (10 µL) were injected onto a Thermo Hypersil Gold C18 column (2.1 mm x 100 mm; 1.9 μm) maintained at 50°C. Mobile phase A consisted of water containing 10 mM ammonium formate and 0.1% (v/v) formic acid. Mobile phase B consisted of a 90/10 (v/v) mixture of isopropanol:acetonitrile containing 10 mM ammonium formate and 0.1% (v/v) formic acid. The LC gradient, maintained at a flow rate of 400 μL/min, was as follows; a starting condition of 65% mobile phase A, 35% mobile phase B. Mobile phase B increased from 35 to 65% over 4 min, followed by 65% to 100% over 15 min, with a hold for 2 min before re-equilibration to the starting conditions over 6 min. All samples were analysed in positive and negative ionisation modes over the mass-to-charge ratio (m/z) range of 250 to 2,000 at a resolution of 100,000. The data was processed with Progenesis QI v2.4 software (Non-linear Dynamics). Relative fold quantification was performed using all ion normalization, with outputs limited to ion signals with intensities that showed 1.5-fold (or greater) difference between the study conditions and displayed statistical significance (p≤0.05). This approach was performed for data acquired in both positive and negative ionization modes. Significant features were then identified using both the Lipid Maps and HMDB databases with a mass error tolerance of 5 ppm. Confident annotations were allocated based on expected adducts for each lipid class versus the detected adducts.

For further network analysis of lipid classes, we used LINEX^2^, a lipid network explorer that integrates lipid-metabolic reactions from public databases to generate a dataset-specific lipid interaction network, revealing candidate enzymes involved in these networks^27^. For this analysis, we used all confident lipid IDs (i.e., >1.5-fold change) from the lipidomics analysis (**Supplementary data 1**). Lipid data files and sample data files with the group names were uploaded on the LINEX^2^ software. The lipid IDs were converted to LipidLynxX Nomenclature and checked against inbuilt *Danio rerio* Rhea and Reactome Databases. The *p* value was set to 0.05 to visualise fold change between lipid overload and RAPA-treated SHK-1 samples.

### Proteomics

Cells were washed and resuspended in an extraction buffer containing 5% SDS, 50mM Triethyl Ammonium Bicarbonate (TEAB), pH 8.5 at sample to buffer ratio of 1:10 (w/v) and homogenized using Precellys homogenizer at 5,000g for 20 sec in a ceramic beads vial (Precellys Lysing Kit, Tissue homogenizing CK mix). The extracts were centrifuged for 10 mins at 16,000g and the supernatant sonicated for 10 cycles (30 sec on, 30 sec off per cycle) on a Bioruptor Pico Sonicator (Diagenode). After sonication, samples were centrifuged (16,000xg for 10 mins), before the supernatant was collected and a bicinchoninic acid assay was performed to assess protein concentration.

The protein samples were digested on the S-Trap micro column (Protifi, USA) following the manufacturer’s protocol with minor modifications. Briefly, 20μg protein in extraction buffer was reduced and alkylated using 10mM dithiothreitol and 40mM iodoacetamide respectively, at 45°C for 15 m. Alkylation was stopped by adding phosphoric acid to a final concentration of 2.5%. The protein solution was diluted six-fold with binding buffer (90% v/v, Methanol in 100 mM TEAB), vortexed gently and loaded into an S-Trap micro column. After centrifugation at 4,000g for 1 m, the column was washed three times with 150μL binding buffer. Proteolytic digestion was carried out by adding 20μL of digestion buffer containing 1μg trypsin in 50 mM TEAB. The column was incubated at 47°C for 2 hr. Digested peptides were eluted using 4 μL of three buffers consecutively: (i) 50mM TEAB, (ii) 0.2% (v/v) Formic acid in H2O, and (iii) 50% (v/v) acetonitrile. Eluted peptides were pooled and cleaned up using C18 stage tips and dried under vacuum.

Purified peptides were separated over a 70 m gradient on an Aurora-25 cm column (IonOpticks Australia) using an UltiMate RSLCnano LC System (ThermoFisher Scientific) coupled to a timsTOF FleX mass spectrometer via a Captivespray ionization source. The gradient was delivered at a flow rate of 200 nL/mins and the column temperature was set at 50°C. The TIMS elution voltage was linearly calibrated to obtain 1/K0 ratios using three ions from the ESI-L Tuning Mix (Agilent, Santa Clara, CA, USA) (m/z 622, 922, 1222) using timsControl (Bruker). For Data Dependent Acquisition (DDA) - Parallel Accumulation-Serial Fragmentation (PASEF) acquisition, the full scans were recorded from 100 to 1700 m/z spanning from 1.45 to 0.65 Vs/cm2 in the mobility (1/K0) dimension. Up to 10 PASEF MS/MS frames were performed on ion-mobility separated precursors, excluding singly charged ions which are fully segregated in the mobility dimension, with a threshold and target intensity of 1750 and 14,500 counts, respectively. The collision energy was ramped linearly from 59 eV at 1/k0 = 1.6 to 20 eV at 1/k0 = 0.6.

Raw mass spectral data was processed using PEAKS Studio X-Pro Software (Bioinformatics LTD)^28^, with protein searches conducted against the Ensembl 113 annotation for Atlantic salmon (https://www.ensembl.org/Salmo_salar/Info/Index), which contains 146,172 unique amino acid sequences (derived from 47,205 coding genes). MS1 precursor mass tolerance was set to 20 ppm, and the MS2 fragment ion tolerance was 0.06 Da. The search parameters specified fully tryptic digestion, allowing for one missed cleavage. Cysteine was treated as a fixed modification with a mass addition of [+57.02], while methionine oxidation and deamination of asparagine and glutamine were set as variable modifications. Quantitative LFQ analysis was performed using default settings, with optional identification transfer enabled.

### Quantification of confocal images using Fiji

The confocal images in Figures 1B, 3A, 5A and 7A were analysed quantitatively using ImageJ-win64 (Fiji app)^25^. For LD size, the diameter of each LD was measured with the freehand line tool. For LD counts and ASM in the extracellular matrix, the multipoint tool was used to count LDs and ASM within each cell and across the entire microscopic field. Colocalization of Plin3 and BODIPY was assessed with the Colocalisation Finder plugin, enabling precise analysis of overlapping signals.

### Statistics and reproducibility

Data are presented as mean ± SD in bar graphs. Statistical significance was determined using two-tailed Student’s t-test via GraphPad Prism software (Version 8.1.1), with *p* values indicated in the corresponding figures. *p* < 0.05 was considered statistically significant for most of the experiments except for proteomics analysis where *p* < 0.01 (adjusted) was considered statistically significant. The exact sample size (*n*) for each experiment is presented in the figure legends. All experiments were performed at least three times independently, yielding consistent results. Statistical analyses from confocal microscopy excluded dead cells showing abnormal nuclear staining, i.e., nuclear condensation.

## Results

### Lipid overload prevents LD breakdown by blocking autophagic flux in SHK-1 cells

We first tested whether LDs generated by lipid overload in SHK-1 cells are regulated by autophagy across a time course, taking samples at 24, 48 and 72 hr following treatment with a 2.5% lipid mixture. To enhance the autophagic flux we added 1μM RAPA^29^ and to block autophagosome formation/maturation we treated the cells with 500nm MRT68921 (MRT)^30^. Firstly, we checked endogenous levels of LD in SHK-1 cells untreated with a lipid mixture, observing negligible BODIPY staining (**Figure 1B**). In comparison, SHK-1 cells treated solely with lipid mixture showed several large circular LDs, mostly in close proximity to the nucleus (**Figure 1B**). Autophagy induction via RAPA treatment resulted in significantly reduced LD number and size (**Figures 1B-1D**). However, treatment with RAPA+MRT led to large LDs at all timepoints, which showed clumping at 72 hr (**Figure 1B**). To avoid clumped LDs that were present at 72 hr, we quantified differences in size and number of LDs across treatment at 48 hr (**Figure 1C-1D**). These results indicate that autophagy induction acts to breakdown LDs, but this effect is ablated by blocking autophagosome formation and maturation.

To further validate the above results, it was important to ask whether LDs colocalise with punctate LC3, the marker for mature autophagosomes, which we achieved by blocking autophagic flux using 10μM Baf-A^31^. Baf-A interrupts autophagic flux by inhibiting V-ATPase-dependent autophagosome-lysosome fusion^32^. This approach aids in visualisation of LD cargo within autolysosomes, the autophagosome-lysosome fusion vesicle. While we imaged SHK-1 cells at all tested time points, LD-LC3 colocalisation was only observed 24 hr post lipid treatment (**Figure 1E**). We specifically observed colocalisation of dense LC3 around large LDs within close proximity to the nucleus (**Figure 1E**). After RAPA treatment, LC3 puncta were observed around several smaller LDs spread across the cytoplasm and also within close proximity of the nucleus (**Figure 1E**). The same colocalisation appeared reduced on the largest LDs following MRT treatment (**Figure 1E**), suggesting LC3 binds to LDs in SHK-1 cells.

To further validate that RAPA was inducing autophagic flux, we measured protein abundance levels of LC3 (with and without addition of Baf-A to all the treatments) and SQSTM1/p62 without addition of Baf-A. LC3 exists as the non-lipidated form LC3-I in the cytoplasm, after lipidation by E1, E2, and E3-like enzymes, respectively (ATG7, ATG3, and a complex consisting of ATG12-ATG5 and ATG16L1), conjugating the cytosolic LC3-I protein to phosphatidylethanolamine to form LC3-II^33^. A combined increase in expression of LC3-II with reduction in SQSTM1/p62 (an autophagy adaptor that gets co-degraded with the cargo) levels is considered to arise from increased autophagic flux^34^. In SHK-1 cells, we observed increased LC3-II expression following RAPA treatment compared to cells solely treated with lipid overload or with additional MRT treatment (**Figures 2A-B**). RAPA treatment reduced SQSTM1/p62 expression, while both lipid overload and MRT treatments enhanced SQSTM1/p62 expression, suggesting a block in autophagic flux on lipid treatment (**Figures 2C-D**). The total protein levels used for normalisation are provided in **Figure 2E**.

**Figure 2.**
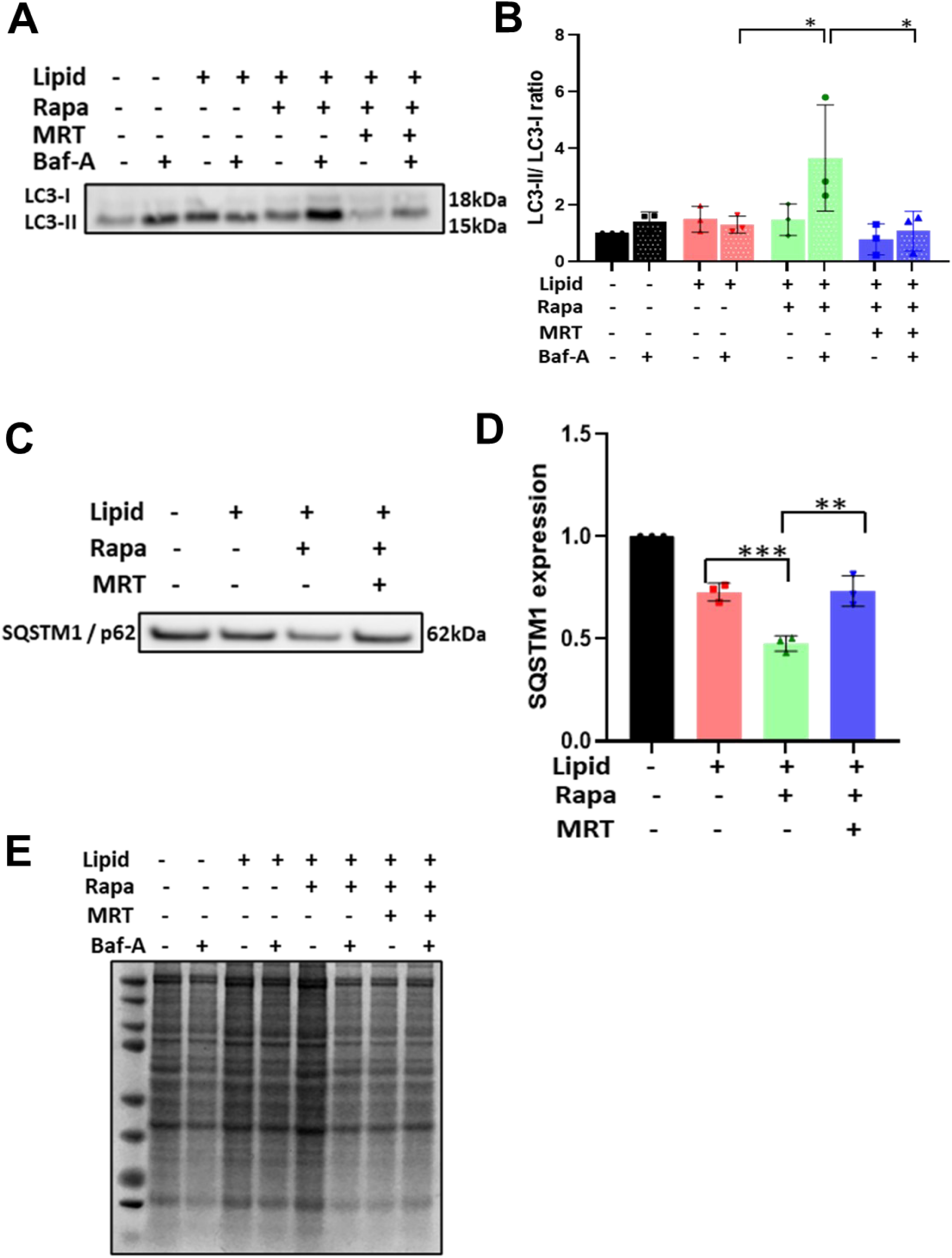
Rapamycin enhances autophagic flux in Atlantic salmon SHK-1 cells under lipid overload. **A.** Representative immunoblot and **B.** Immunoblot quantification showing LC3 abundance in SHK-1 cells after 72 hr treatment with no lipid overload, lipid overload, lipid overload with RAPA, and lipid overload with RAPA and MRT treatments, with and without Baf-A (n=3). **C**. Representative immunoblot and **D.** Immunoblot quantification showing SQSTM1/p62 abundance in SHK-1 cells after 72 hr treatment with no lipid overload, lipid overload, lipid overload with RAPA, and lipid overload with RAPA and MRT treatments. **E.** Total protein blot stained with simply blue safe stain for protein abundance quantification. Data shown as mean +/− SD, * *p* < 0.05; ** *p* < 0.01.

### RAPA enhances lipid droplet breakdown via perilipin 3 and upregulation of key lipases

Our next question was whether LDs in SHK-1 cells colocalise to Plin3, a protein that coats LDs and acts as a key marker for lipophagy. Plin3 is required for docking autophagy machinery to LDs and its silencing blocks LD-associated LC3-II autophagic flux^35^. LD-Plin3 co-localisation was observed at all three time points after blocking autophagic flux with Baf-A. Blockage of autophagic flux led to accumulation of more small LDs in RAPA-treated cells compared to the lipid overload and MRT treated cells, which showed bigger LDs (**Figures 3A-3B**, 72 hr data shown). The small LDs observed in RAPA-treated cells were further surrounded with more Plin3 compared to LDs in lipid overload (**Figures 3A-3B**). Plin3 was strongly associated with small LDs, as assessed by Pearson’s coefficient (**Figure 3C**), with large LDs showing a more limited association with Plin3, especially in cells with lipid overload. These results clearly suggest a role for Plin3 in RAPA-induced LD breakdown in SHK-1 cells.

**Figure 3.**
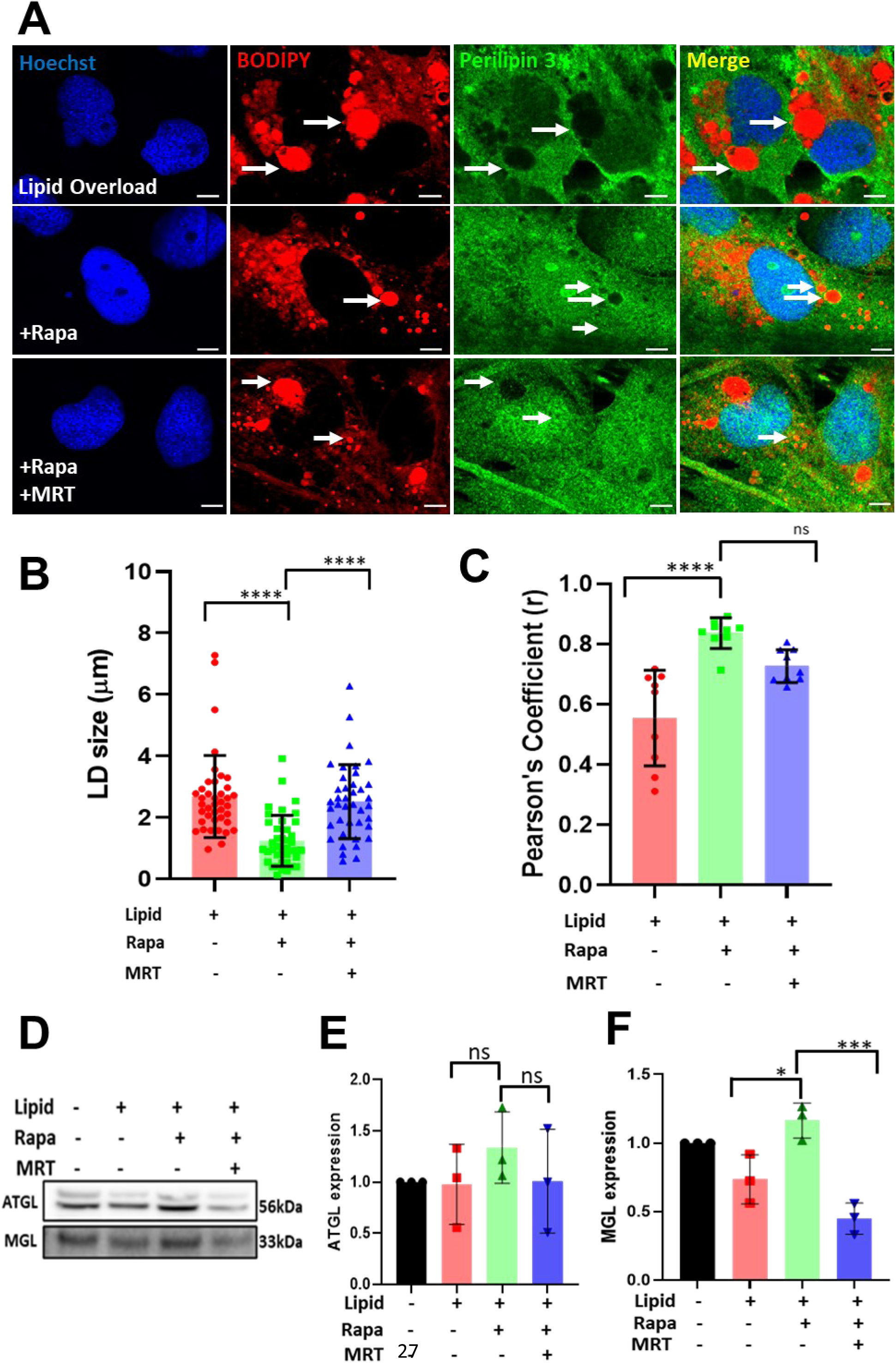
Rapamycin enhances Lipid droplet breakdown via perilipin 3 and upregulation of key lipases. **A**. Representative confocal images showing the co-localisation of Plin3 (green puncta) with LD stained with BOPIPY (red) in SHK-1 cells after 72 hr treatment with lipid overload, lipid overload with RAPA, and lipid overload with RAPA and MRT treatments. Baf-A was added to all treatments to block the autophagic flux. Scale bar = 10μm, n=3. **B.** Quantification of lipid droplet size (40 cells were counted from 3 independent experiments). **C.** Quantification of co-localisation between Plin3 and lipid droplets as measured by Pearson’s correlation co-efficient. **D.** Representative immunoblot and **E-G.** Immunoblot quantification showing ATGL and MGL abundance in SHK-1 cells after 72 hr treatment with no lipid overload, lipid overload, lipid overload with RAPA, and lipid overload with RAPA and MRT treatments. The blots were normalised against total protein in Fig 2E (n=3). Data shown as mean +/− SD, * *p* < 0.05; *** *p* < 0.001; **** p < 0.0001, ns= non-significant.

We further assessed the expression of key lipase proteins, i.e., adipose triglyceride lipase (ATGL)^36^ and monoglyceride lipase (MGL)^37^ in SHK-1 cells treated with lipid overload, RAPA and MRT. The abundance of both the proteins was reduced under both the lipid overload and when the autophagosome formation was blocked with MRT treatment for 72 hr (**Figures 3D-F**). Together, these results suggest that SHK-1 cells have all the key molecules required for LD breakdown via autophagy, which are inhibited by lipid overload, a process that can be experimentally reversed by enhancing autophagic flux.

### Autophagy supresses bioactive ceramides and upregulates PUFA-TAGs

To gain a global perspective on the role of lipid autophagy in SHK-1 cells, we examined changes in lipid pool abundance using lipidomics, following the induction of autophagy with RAPA in comparison to the lipid overload treatment (72 hr post-treatments). The global lipidome of SHK-1 cells under lipid overload exhibited an increased abundance (>2-fold change) of SMs, including SM 36:0 and SM 36:1. In contrast, cells treated with RAPA showed an increased abundance (>2-fold change) of long-chain PUFA-triacylglycerols (TAGs) containing 55-60 carbons with 5-9 double bonds, alongside ion signals likely attributable to TAGs containing polyunsaturated fatty acids (**Figures 4A-B**).

**Figure 4.**
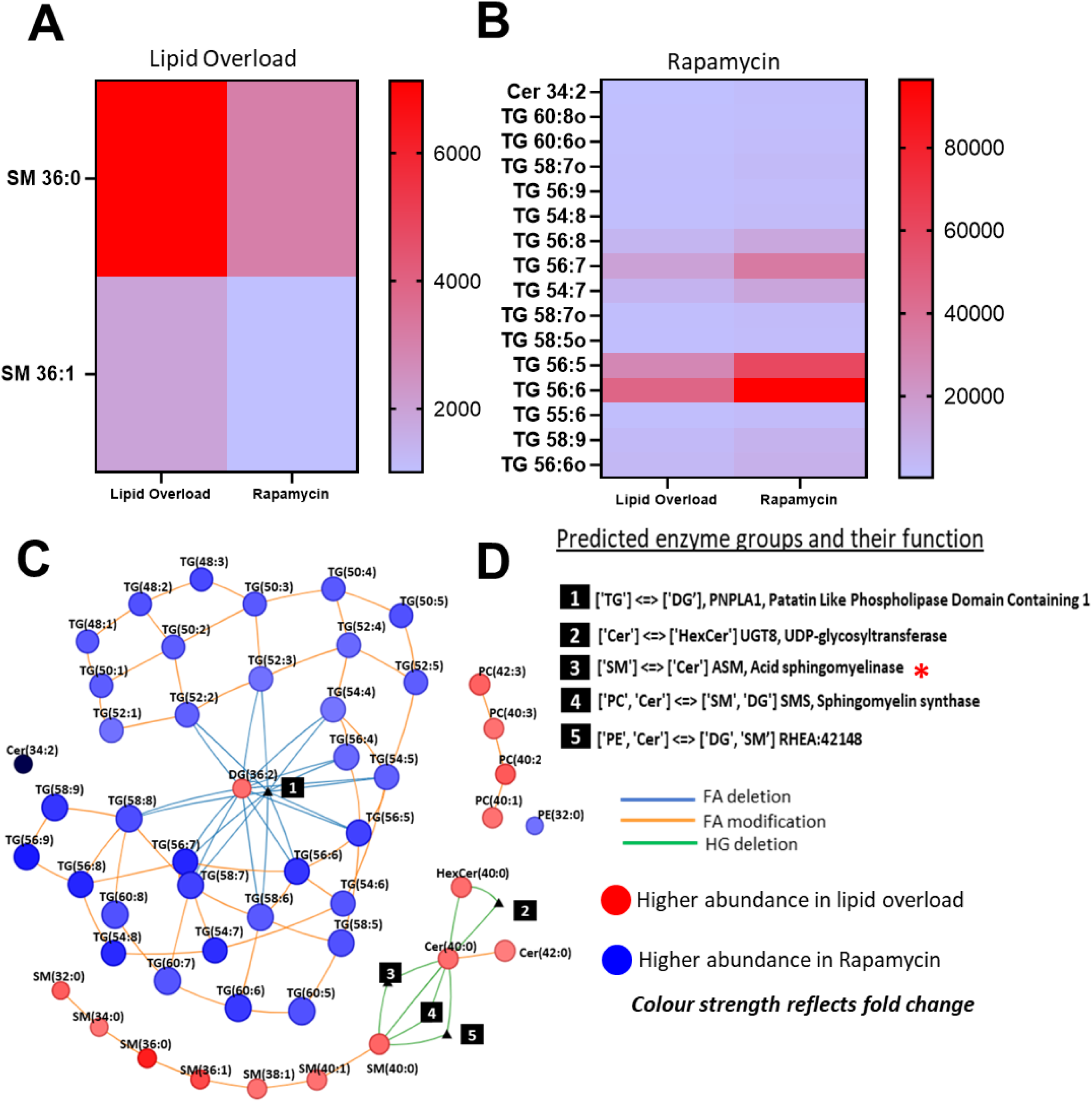
Autophagy supresses bioactive ceramides and upregulates the PUFA-TAG pool. SHK-1 cells treated with lipid overload and lipid overload with RAPA treatment for 72 hr were analysed for lipidome changes. **A.** Heat map of saturated SMs (>2 fold up-regulated) under lipid overload treatment. **B.** Heat map of PUFA-TAGS (>2 fold up-regulated) under lipid overload and RAPA treatments. **C**. LINEX^2^ network enrichment analysis showing two key networks, the TAG network upregulated under RAPA treatment (blue circles) and ceramide-sphingomyelin network upregulated under lipid overload treatment (red circle) (both including lipid species showing >1.5-fold change). **D.** The embedded table shows five enzymes responsible for the networks shown were predicted by LINEX^2^ based on metabolite abundances of SHK-1 cells following lipid overload and lipid overload with RAPA. The red asterisk indicates acid sphingomyelinase, which was further investigated. Data shown *p* < 0.05.

Lipotoxicity is associated with altered levels of CER, diacylglycerols, and SM^38–40^. SM 36:0 and SM 36:1, identified under lipid overload, have been shown to be upregulated in lipid metabolic disorders such as fatty liver, liver steatosis in NAFLD, high-fat diets, and obesity^39,41–43^. To further explore evidence for lipotoxicity in our data, we employed LINEX^2^, a lipid network explorer that integrates lipid-metabolic reactions from public databases to generate a dataset-specific lipid interaction network (**Figure 4C**), revealing candidate enzymes which may be involved in these networks^27^. The LINEX^2^ network enrichment revealed two key networks, the TAG network upregulated following RAPA treatment, and the CER-SM network upregulated by the lipid overload treatment (**Figure 4C**). LINEX^2^ identified five candidate enzymes based on metabolite abundances between lipid overload and SHK-1 cells treated with RAPA (**Figures 4C-D**). Among the predicted enzymes for the lipid overload treatment, we selected acid sphingomyelinase (ASM), which resides in lysosomes^44^and it is likely to impact or be impacted by blocked autophagic flux seen in lipid overload treatments^45,46^ (**Figure 4D**), for further analysis.

ASM is mainly a lysosomal lipid hydrolase, where it cleaves the sphingolipid, sphingomyelin, into CER and phosphocholine. Knockout of ASM in hepatocytes inhibits palmitic acid induced lipotoxicity^47^. ASM is also known to localise to the plasma membrane and the extracellular space^44^. We therefore asked whether lipid overload or RAPA treatment alters the localization of ASM, and if this is dependent on autophagy. We performed immunofluorescence staining for ASM and the key lysosomal marker LAMP-1 (lysosomal-associated membrane protein 1) after treating SHK-1 cells with a lipid mixture, the same lipid mixture plus RAPA, and after subsequently blocking autophagy with MRT (**Figure 1A**).

Under lipid overload, ASM predominantly localized within the lysosomes (**Figure 5A**). However, treatment with RAPA induced the exocytosis of ASM into the extracellular space via LAMP-1 positive vesicles (**Figures 5A, white arrows**). Interestingly, when autophagy was blocked using MRT, we observed an increase in the retention of ASM within LAMP-1 positive lysosomes, particularly around the nucleus (**Figure 5A**). These findings suggest that in the absence of autophagy, ASM remains confined to lysosomes, highlighting autophagy’s essential role in regulating its localization (**Figure 5B)**. As an alternative approach, we assessed SMase activity using a fluorimetric sphingomyelinase assay in SHK-1 cells under the same experimental conditions. RAPA treatment resulted in reduced SMase activity compared to cells treated with the lipid mixture alone, while SMase activity significantly increased when autophagy was blocked with MRT (**Figure 5C**). These results indicate a role for autophagy in modulating SMase activity.

**Figure 5.**
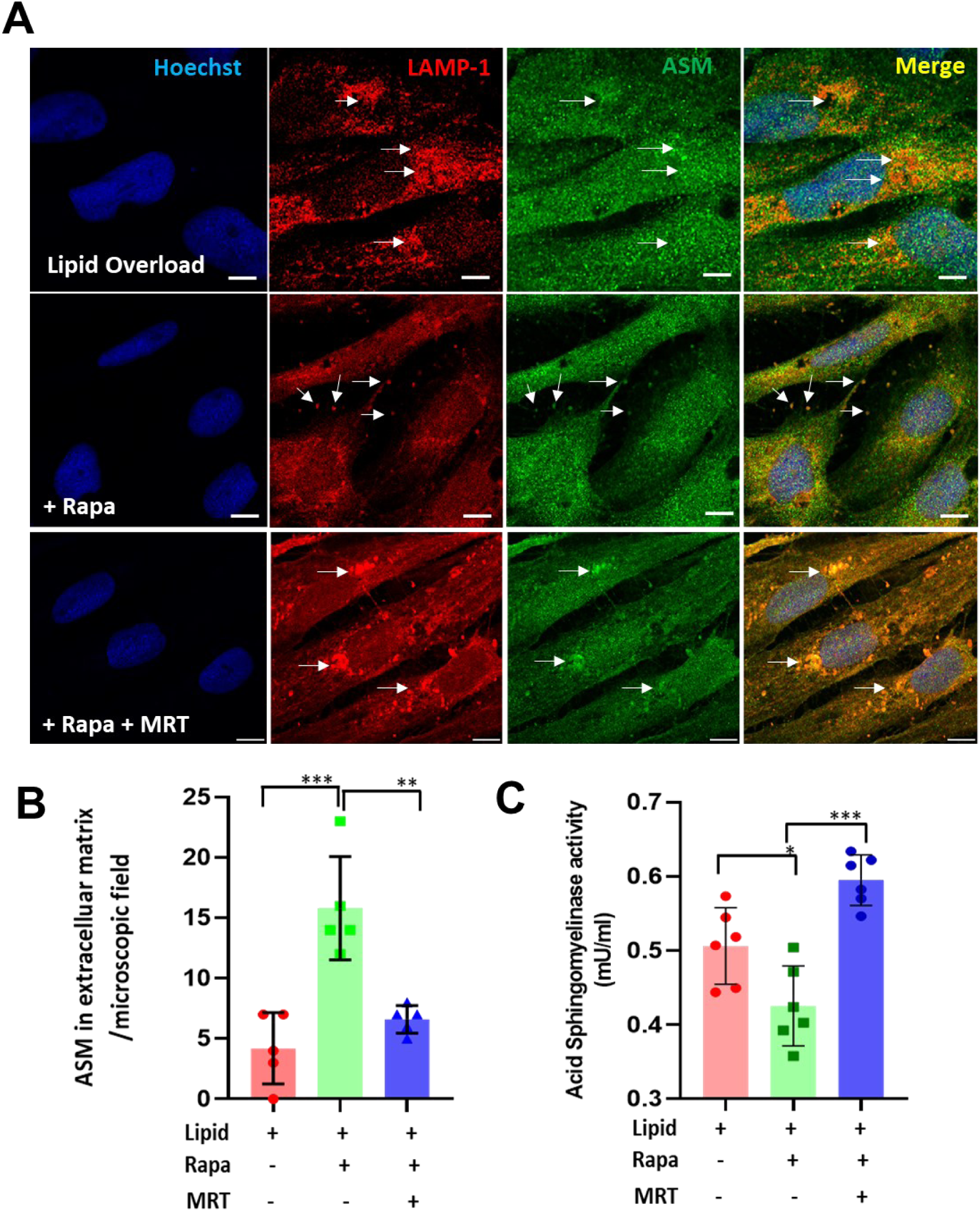
Autophagy regulates ASM localization and modulates SMase activity. **A.** Representative confocal images showing the localisation of ASM (green) with LAMP-1 (red) in SHK-1 cells following 72 hr treatment with lipid overload, lipid overload with RAPA and lipid overload with RAPA and MRT treatments. Scale bar = 5μm, n=3. **B.** Quantification of extracellular localisation of ASM with LAMP-1 positive vesicles (5-6 microscopic fields with minimum 9-10 cells each were used from 3 independent experiments). **C.** Evaluation of Acid SMase activity using fluorimetric assay (n=6). Data shown are mean +/− SD, * *p* < 0.05; ** p < 0.01; *** *p* < 0.001

### Rapamycin ameliorates lipid overload mediated lipotoxicity in SHK-1 cells

To better understand the functional impacts of autophagy induction on SHK-1 cells, we performed a proteomics analysis comparing the lipid overload vs. RAPA treatments after 72 hr (**Figure 1A**). 5,971 proteins were confidently detected in SHK-1 cells (**Supplementary data 2**). We focused on those showing changes in abundance > 1.5-fold increase or < 0.5-fold decrease (*p* <0.01) (**Figure 6A**). This approach identified 20 upregulated and 3 downregulated proteins under lipid overload. Among the 21 up-regulated proteins, one was uncharacterized. 13 of the remaining 20 Atlantic salmon proteins were orthologues of human proteins associated with fatty liver disease according to a published database^48^ (**Figure 6B**), along with 2 of the 3 downregulated proteins, suggesting a strong level of functional conservation across distant vertebrate species. These 16 proteins were arranged in descending order of their standardized value scores (log10 p-value) to represent the relative strength of functional association with fatty liver pathology in human (**Figure 6C**).

**Figure 6.**
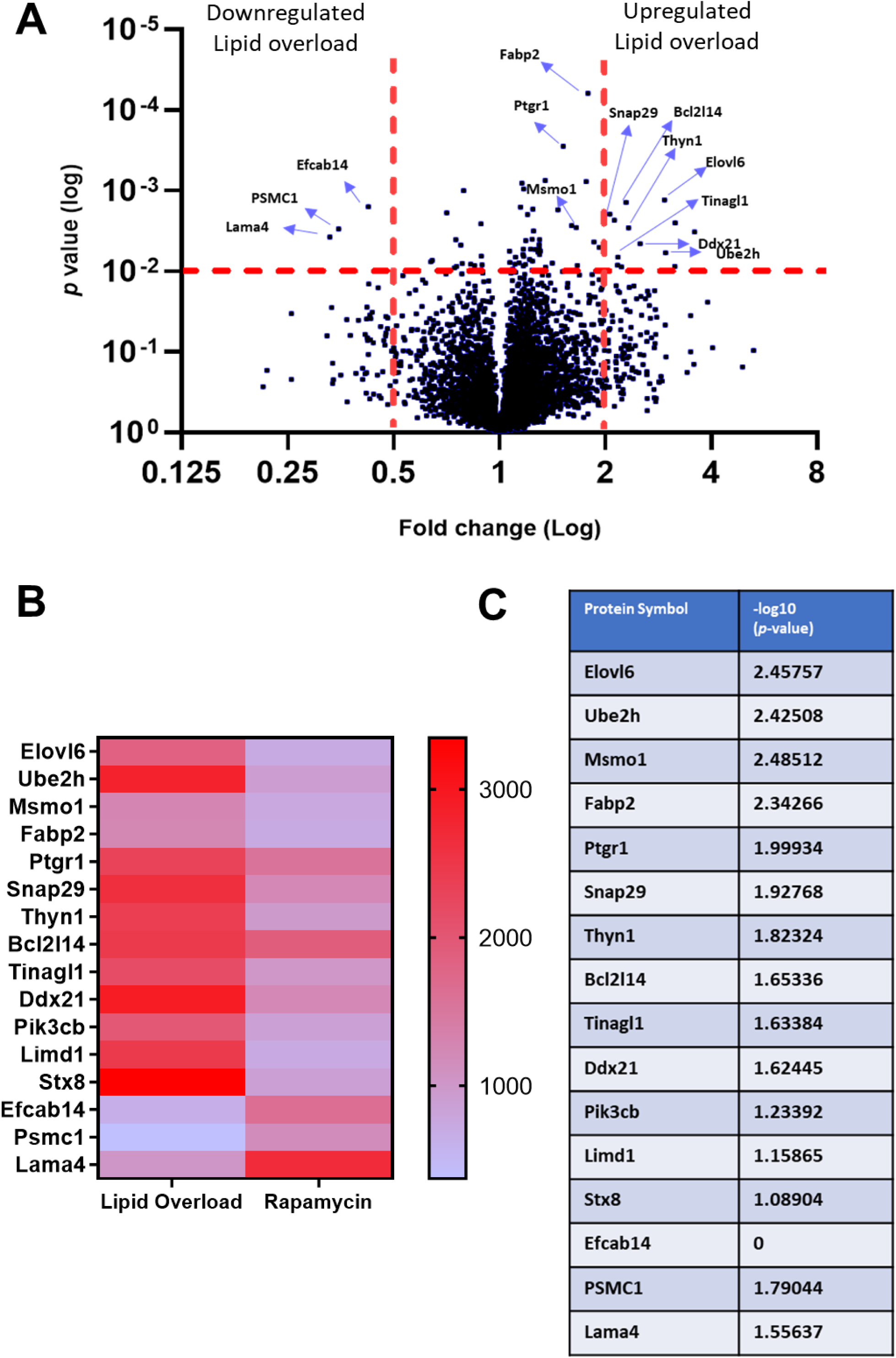
Rapamycin ameliorates lipid overload mediated lipotoxicity. SHK-1 cells with lipid overload or lipid overload with RAPA treatment were analysed for proteome changes (72 hr post-treatments). **A.** Volcano plot highlighting key protein targets among total 5,971 proteins identified. **B.** Heat map of proteins showing abundance changes in the lipid overload treatment with < 0.5-fold change or >2-fold change (*p <* 0.01). **C**. Standardized value scores (log10 p-value) to represent the relative strength of the functional association of key protein targets with fatty liver (scores obtained from CTD Gene-Disease Associations dataset ^77^).

### ELOVL6 as a target for selective autophagy in SHK-1 cells

Among the 16 proteins upregulated by lipid overload, we selected ELOVL6 (fatty acid elongase 6) for additional validation and downstream analysis, as it had the highest functional association score with fatty liver pathology (**Figure 6C**). ELOVL6 is involved in the elongation of saturated and monounsaturated fatty acids, particularly converting 12, 14 and 16 carbon fatty acids^49^. Its role in mammalian lipid metabolism has drawn significant attention in the context of metabolic diseases such as NAFLD and obesity due to its influence on lipid accumulation, inflammation, and insulin resistance^50^. Inhibiting ELOVL6 activity may help prevent the conversion of palmitate to longer-chain fatty acids, thus reducing lipotoxicity and inflammasome activation, which are crucial factors in the progression of fatty liver disease^50^. We analysed changes in ELOVL6 protein levels in SHK-1 cells using immunofluorescence. Our results showed an increase in ELOVL6 abundance in cells treated with lipid overload and when autophagy was blocked using MRT, compared to those treated with RAPA (**Figures 7A-B**). When autophagic flux was blocked with MRT and Baf-A, we observed a significant aggregation of ELOVL6 in SHK-1 cells treated with lipid overload, indicating an accumulation of ELOVL6 protein in the absence of autophagy (**Figures 7A-C**).

**Figure 7.**
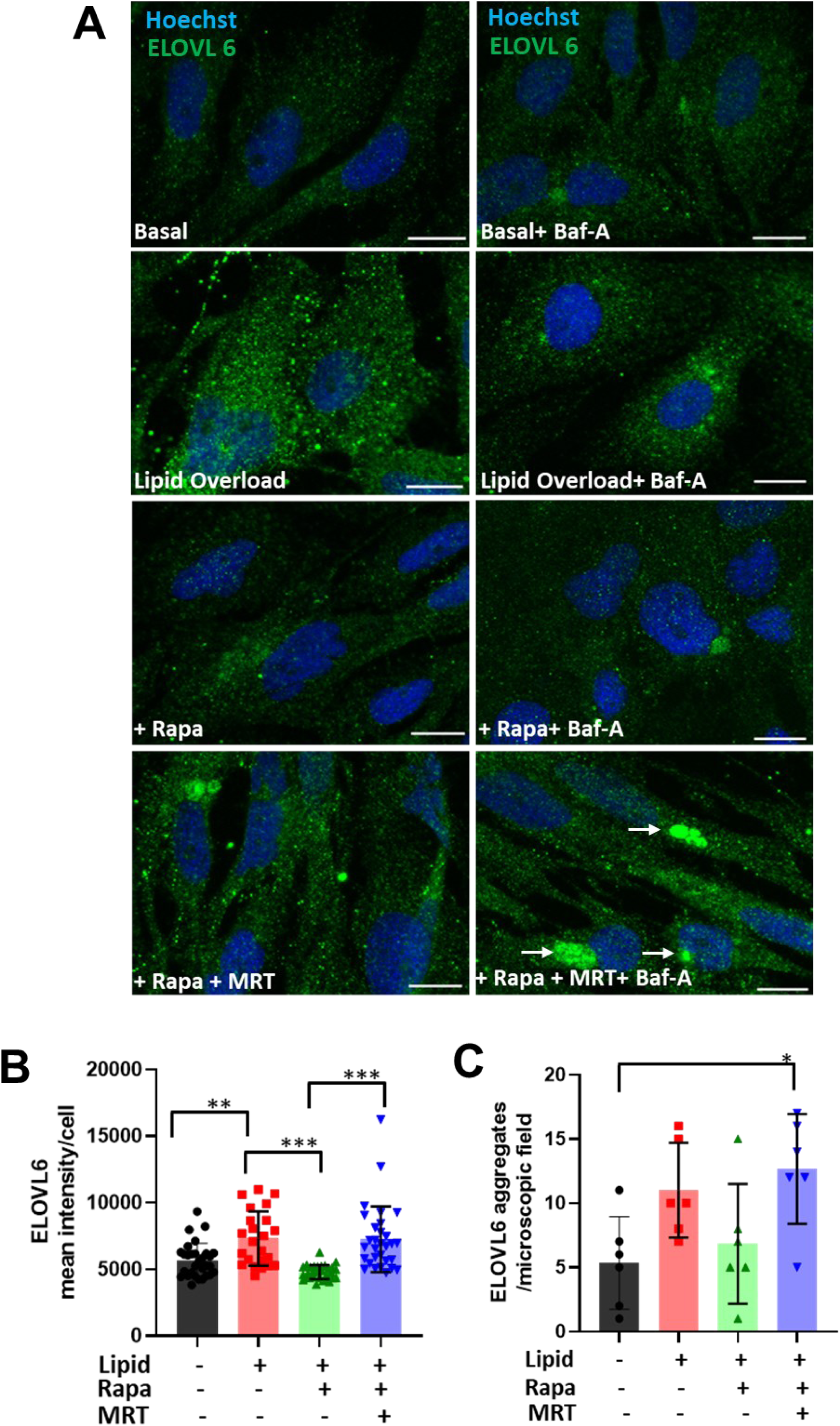
ELOVL6 is a possible target for autophagosomes. Representative confocal images showing expression of ELVOL6 in SHK-1 cells after 72 hr treatment with no lipid overload, lipid overload, lipid overload with RAPA, and lipid overload with RAPA and MRT treatments with and without Baf-A treatment to block autophagic flux. **A-B.** Increased expression of ELOVL6 in lipid treated cells and its quantification (3 microscopic fields with minimum 9-10 cells each were used from 3 independent experiments). **A and C** Increased aggregation of ELOVL6 on inhibition of autophagic flux with MRT and Baf-A (3 microscopic fields with minimum 9-10 cells each were used from 3 independent experiments). Scale bar = 10μm, n=3. White arrows indicate ELOVL6 aggregations.

Based on these findings, we hypothesized that ELOVL6 is a target for autophagosomes. To provide further evidence of its targeting by autophagy, we asked whether Atlantic salmon ELOVL6 proteins have a LC3-interacting region motif (LIR motif). The LIR motif is a short linear sequence that facilitates the interaction between autophagy receptors and LC3 or other ATG8 family proteins^51^. The canonical LIR motif is characterized by a specific sequence pattern: Θ0-X1-X2-Γ3, where: ‘Θ’ (Theta) represents an aromatic residue, which can be tryptophan (W), phenylalanine (F), or tyrosine (Y). This residue is critical for the interaction with the hydrophobic pocket of LC3 or ATG8 proteins. ‘X’ represents any amino acid (aa), ‘Γ’ (Gamma) denotes an aliphatic residue, which can be leucine (L), valine (V), or isoleucine (I). This residue helps stabilize the interaction by binding to a different hydrophobic pocket of the LIR docking site. The arrangement of these residues enables the LIR motif to bind to the hydrophobic pockets of LC3 or ATG8 family proteins, promoting the recruitment of autophagy receptors to sites of autophagosome formation and facilitating autophagic degradation of the target protein^52^.

The Atlantic salmon genome contains many co-orthologues of mammalian genes, owing to a history of fish-specific whole genome duplication events during evolution^12^. Consistently, we identified five highly similar *elovl6* paralogues annotated in the NCBI genome (Ssal_v3.1, GCA_905237065.2), each encoding products sharing extensive homology with human ELOVL6. All five Atlantic salmon proteins were checked for a LIR motif using the iLIR Autophagy database^51^ (**Supplementary data 3**). Specifically, each protein showed at least one xLIR motif (PSSM score of 10 or 11) and several WxxL motifs (PSSM score ranging from 1-9). One of the paralogues (XP_014034459) showed evidence for two xLIR motifs with a PSSM score of 10 and 11 (p < 0.001) and several WxxL motifs (**Supplementary data 3**). None of these motifs were present in the anchor region. These findings suggest that Atlantic salmon ELOV6 likely contains the expected motif to be targeted by selective autophagy.

## Discussion

Lipid recycling by autophagy, also known as lipophagy, has been extensively studied in mammals, where it plays crucial roles in LD turnover in hepatocytes^53^ and adipocytes^54^. In *Drosophila*, which shares its core lipid metabolism pathways with vertebrates, small GTPase Rab40 is required for mobilisation of lipids via lipophagy^55^. Furthermore, RAPA treatment enhances lipophagy to prevents photoreceptor degeneration by blocking the formation of dihydroceramide, a precursor to CER accumulation in the *Drosophila* eye^56^. Lipophagy also plays important roles in lipid metabolism in zebrafish (*Danio rerio*)^3,57^, where the simultaneous inhibition of cytosolic lipolysis and lipophagy resulted in severe liver damage with inflammation and damaged mitochondria^58^. Until this study, the role of lipophagy in Atlantic salmon has remained uncharacterised, an important knowledge gap considering the potentially key role that lipid homeostasis plays in the health of farmed fish. Using SHK-1 cells, we established a lipid overload model to investigate both lipotoxicity and lipid recycling by autophagy. This *in vitro* system offers flexibility for testing various autophagy inducers and inhibitors in a time and dose-dependent manner, while avoiding the use of live fish - aligning with principles of the 3Rs (Replacement, Reduction, and Refinement) in research^59^

Our experiments demonstrate reduced autophagic flux in Atlantic salmon SHK-1 cells under lipid overload, as indicated by the increased expression of SQSTM-1/p62 (**Figure 2**). Additionally, a lipid overload treatment led to no significant change in LC3-I to LC3-II conversion following Baf-A treatment, suggesting a blockage in autophagic flux (**Figure 2**). This parallels findings in vascular smooth muscle cells treated with saturated fatty acids, where a similar block in autophagic flux contributes to lipotoxicity^60^. We also observed accumulation of large LDs lacking Plin3 co-localisation and reduced protein levels of the three key lipase enzymes under lipid overload (**Figure 3)**, further suggesting inhibition of LD breakdown both by autophagy and cytosolic lipases.

Lipidomics under lipid overload conditions revealed an up-regulation of several saturated SMs and CERs (**Figure 4**), the key lipotoxic molecules implicated in various metabolic disorders^22,61^. *De novo* CER synthesis, fuelled by saturated fatty acids like palmitic acid^62^ contributes to lipotoxicity by accumulating in LDs^63^. Additionally, SMs are converted to CERs by ASMs^64^. Our analysis pinpointed ASM as a key enzyme that co-localized with lysosomes under lipid overload, suggesting it represented lysosomal ASM, with a concomitant increase in ASM activity (**Figure 5**). Proteomics also showed an upregulation (>2-fold) of proteins associated with fatty liver; a condition manifested by lipotoxicity (**Figure 6**). Importantly, our experiments reveal that this lipotoxicity is effectively suppressed by enhancing autophagy. RAPA treatment not only reduced the accumulation of lipotoxic molecules, but also restored autophagic flux, evidenced by decreased SQSTM-1/p62 levels, enhanced LC3-I to LC3-II conversion, and a reduction in the number and size of LDs. Reduction in the number of LDs by RAPA treatment has elsewhere been demonstrated in palmitic acid induced foam cells^65^.

Furthermore, RAPA treatment resulted in the accumulation of highly unsaturated TAGs, as indicated by the presence of multiple double bonds, which have a protective role against lipotoxicity induced by saturated fatty acids^66,67^. Based on the high number of double bonds observed in the high-carbon TAGs, we hypothesize that these TAGs are enriched in omega-3 fatty acids, which will require further experiments to test. Within the bounds of this study, attempts at measuring omega-3 levels using ELISA were unsuccessful, presumably due to the very low levels of DHA in SHK-1 cells (data not shown). This highlights the need for alternative methods or optimization of the assay to accurately measure omega-3 fatty acids in SHK-1 cells. Interestingly, RAPA-induced accumulation of TAGs has also been reported in microalgae, albeit with limited information on carbon chain length and the number of double bonds involved^68^. Enhanced accumulation of PUFA-TAGs by RAPA is possibly a protective mechanism against toxicity caused by reactive species produced by oxidation of membranous TAGs and associated increases in reactive oxygen species^67^.

It is also worth noting that RAPA treatment resulted in exocytosis of ASM in LAMP-1 positive vesicles if SHK-1 cells. This secretory form of ASM plays a role in plasma membrane repair when cells are wounded in the presence of elevated calcium ions ^69,70^. While, we did not measure calcium ion concentration in our treatments, enhanced calcium levels have been reported in the serum of patients with dysregulated lipid profiles ^71^.

By blocking the formation of autophagosomes using MRT, we further strengthened our claim that the suppression of lipotoxicity is indeed autophagy-dependent in SHK-1 cells. Blocking autophagic flux with MRT resulted in a significant increase in LD size, ASM localisation to the lysosomes and an increase in ELOVL6 aggregation, supporting the hypothesis that autophagy is required to regulate the lipotoxic environment. This further suggests that the autophagy pathway is crucial in mitigating lipotoxic effects by managing lipid storage and degradation processes.

In humans and mice, autophagy levels decline with age, which also increases susceptibility to lipotoxicity^72–76^. However, such data is not yet available for farmed Atlantic salmon. Our findings suggest that enhancing autophagy in salmon cells can restore lipid homeostasis, presenting a promising approach to counteracting lipotoxicity induced by dietary factors (i.e., increased vegetable oils in farmed fish diets) or a sedentary domesticated lifestyle, particularly as Atlantic salmon age during the course of aquaculture production. In this respect, using dietary supplementation to activate autophagy may offer a novel strategy to produce healthier fish with improved lipid profiles, which may lead to increased robustness. As an obvious next step, our *in vitro* results should be validated using alternative autophagy inducers, especially those amenable for use as *in vivo* feed supplements on large fish during the final stage of production leading to harvest.

## Acknowledgements

We thank Dr Anna Raper (Bioimaging Facility Manager, Roslin Institute) for support with the immunofluorescence experiments. We thank, Dr Ye Hwa Jin and Dr Diego Robledo from The Roslin Institute for their help with SHK-1 cell culture.

## Funding Sources

This work was supported by funding from the Scottish Universities Life Sciences Alliance (SULSA ECR Development funding to K.P.), the BBSRC (Roslin Institute pump priming grant to K.P. and Institute Strategic Programme grants BB/J004316/1; BBS/E/RL/230001C and BBS/E/D/10002071) and the Seafood innovation fund (grant RD204 to K.P.).

## Abbreviations

Lipid droplets (LDs), ceramides (CERs), sphingomyelins (SMs), salmon head kidney cell line (SHK-1), Rapamycin (RAPA), MRT68921 (MRT), microtubule-associated light chain 3 (LC3), Perilipin 3 (Plin3), triacylglycerol (TAG), fatty acid elongase 6 (ELOVL6) and acid sphingomyelinase (ASM), Standard Deviation (SD)

## Supplementary Data

**Supplementary Data 1.** Complete lipidomics dataset both in the positive ion mode and the negative ion mode from SHK-1 cells treated with lipid overload and lipid overload and RAPA.

**Supplementary Data 2.** Complete data and list of differentially regulated proteins in SHK-1 cells treated with lipid overload and lipid overload and RAPA.

**Supplementary Data 3**. Presence of xLIR and Wxxl motifs in five *elovl6* paralogues identified in the Atlantic salmon genome.

## Notes

### Competing Interest Statement

The authors have declared no competing interest.

