## Supplementary Data 3 for "Rapamycin induced autophagy enhances lipid breakdown and ameliorates lipotoxicity in Atlantic salmon cells"

**Supplementary data 3: Five paralogues of ELOVL6 in Salmo salar with putative LIR motifs**

1. **Elongation of very long chain fatty acids protein 6 [Salmo salar]**

NCBI Reference Sequence: XP_014054664.1

>XP_014054664.1 elongation of very long chain fatty acids protein 6 [Salmo salar]

MSVLALQEYEFERQFNEDEAIRWMQENWKKSFLFSALYAACILGGRHVMKQREKFNLRKPLVLWSLTLAVFSIFGAVRTGSYMTYILMTKGLKQSVCDQSFYNGPVSKFWAYAFVLSKAPELGDTLFIVLRKQKLIFLHW

YHHITVLLYSWYSYKDMVAGGGWFMTMNYLVHSVMYSYYALRAAGFKVSRKFAMFITLTQITQMLVGCVVNYLVYSWMQQGQECPSHVQNIVWSSLMYLSYFVLFCQFFHEAYIDKTKKAAAAKAEAAAAAARKTQ

**Motif	Start	End	Pattern	PSSM Score**

**xLIR	193	198	AMFITL	11**

WxxL	32	37	FLFSAL	5

WxxL	72	77	SIFGAV	2

WxxL	110	115	WAYAFV	2

WxxL	125	130	TLFIVL	6

WxxL	139	144	HWYHHI	1

WxxL	176	181	YSYYAL	6

**WxxL	231	236	IVWSSL	9**

**WxxL	239	244	LSYFVL	8**

### 2. Elongation of very long chain fatty acids protein 6 [Salmo salar]

NCBI Reference Sequence: XP_014067429.1

>XP_014067429.1 elongation of very long chain fatty acids protein 6 [Salmo salar]

MSVLALQEYEFEKQFNEDEAIRWMQENWKKSFLFSAFYAAFILGGRRVMKRREKFELRKPLVLWSLTLAV

FSIFGAVRTGSYMTYILMTKGLKQSVCDQSFYNGPVSKFWAYAFVLSKAPELGDTLFIVLRKQKLIFLHW

YHHMTVLLYSWYSYKDMVAGGGWFMTMNYLVHAVMYSYYALRAVGFKVSRKFAMFITLTQITQMLVGCVVNYLVYSWMQQGQECPSHMQNIVWSSLMYLSYFVLFCQFFHEAYIDKTKEAAAAKTEATAAARKTQ

**Motif	Start	End	Pattern	PSSM Score**

**xLIR	193	198	AMFITL	11**

WxxL	72	77	SIFGAV	2

WxxL	110	115	WAYAFV	2

WxxL	125	130	TLFIVL	6

WxxL	176	181	YSYYAL	6

**WxxL	231	236	IVWSSL	9**

**WxxL	239	244	LSYFVL	8**

**3. Elongation of very long chain fatty acids protein 6 [Salmo salar]**

>XP_014057592.1 elongation of very long chain fatty acids protein 6-like [Salmo salar]

MNESEHQVPLSQYGFEQQFDERGAIEWMQENWSKAFVFCGLYAALVFGGQHFMKERPKLSLRRPLVLWSLSLAIFSIIGAIRTGWYMRYILSTSGFRQSICDQSFYNGPVSKFWAYAFVLSKAPELGDTAFIVLRKQRLI

FLHWYHHITVLLYSWYSYKDQVAGGGWFMTMNYSVHALMYSYYAVRAAGYRVPRPFAVVITTFQIAQMAMGLTVSWLVYHWMQEGDCNSYLNNIVWSSLMYLSYLVLFSSFFYQTYMKGKGGKSSKMD

**Motif	Start	End	Pattern	PSSM Score**

**xLIR	242	247	LSYLVL	11**

WxxL	36	41	FVFCGL	1

WxxL	73	78	AIFSII	5

WxxL	114	119	WAYAFV	2

WxxL	129	134	TAFIVL	7

WxxL	143	148	HWYHHI	1

WxxL	180	185	YSYYAV	5

WxxL	194	199	RPFAVV	5

**WxxL	234	239	IVWSSL	9**

### 4. Elongation of very long chain fatty acids protein 6-like [Salmo salar]

NCBI Reference Sequence: XP_014014538.1

 >XP_014014538.1 elongation of very long chain fatty acids protein 6-like [Salmo salar]

MNETENIPLAEYDFERQFDEKGALEWMQENWSKSFMFCGLYAAIIFGGQHFMRERPKLNLRRPLVLWSLSLAIFSIIGAVRTGWYMFHVLNKSGFKQSVCDTSFYSAPVSKFWAYAFVLSKAPELGDTVFIVLRKQRLIF

LHWYHHITVLVYSWYSYKDQVAGGGWFMTMNYLVHSFMYTYYAARAAGYRVPRPCAMVITATQIMQMMMGLAVLGLVYHWMHEVRCPSYMPNIAWGSLMYLSYLVLFASFFYKSYLKGAAYKVGKGSKAE

**Motif	Start	End	Pattern	PSSM Score**

**xLIR	241	246	LSYLVL	11**

WxxL	35	40	FMFCGL	5

WxxL	72	77	AIFSII	5

WxxL	85	90	YMFHVL	6

WxxL	113	118	WAYAFV	2

WxxL	128	133	TVFIVL	6

WxxL	142	147	HWYHHI	1

**WxxL	233	238	IAWGSL	8**

### 5. Elongation of very long chain fatty acids protein 6-like [Salmo salar]

NCBI Reference Sequence: XP_014034459.1

>XP_014034459.1 elongation of very long chain fatty acids protein 6-like [Salmo salar]

MNETEHIPLAEYDFERKFDEKGALEWMQENWSKSFMFCGLYAAIIFGVQHFMRERPKLNLRRPLVLWSLSLAIFSIIGAVRTGCYMFHVLNKSGFKQSVCDTSFYSAPVSKFWAYAFVLSKAPELGDTVFIVLRKQRLIF

LHWYHHITVLVYSWYSYKDQVAGGGWFMTMNYLVHSFMYTYYAARAAGYRVPRPCAMVITATQIMQMVMGLAVLGLVYHWMHEVRCPSYVPNIAWGSLMYLSYLVLFASFFYKSYLKVAADKESKAK

**Motif	Start	End	Pattern	PSSM Score**

**xLIR	241	246	LSYLVL	11**

**xLIR	253	258	KSYLKV	10**

WxxL	35	40	FMFCGL	5

WxxL	72	77	AIFSII	5

WxxL	85	90	YMFHVL	6

WxxL	113	118	WAYAFV	2

WxxL	128	133	TVFIVL	6

WxxL	142	147	HWYHHI	1

**WxxL	233	238	IAWGSL	8**
